# Plant Breeding in the face of climate change

**DOI:** 10.1101/2022.10.07.511293

**Authors:** Carlos D Messina, Mark Cooper

## Abstract

Climate change will have a net negative and inequitable impact on agriculture. Genetics for crop improvement ranks in the top set of technologies that can contribute to human adaptation to climate change. However, a framework for how to breed crops for climate change adaptation is lacking. Here we propose a framework to develop new genotype (G) x management (M) technologies (G x M) to adapt to climate change, and to transition from current to future G x M technologies in a way that future food security does not come at the expense of current food security. The framework integrate genomic, agronomic, and environmental (E) predictors to accomplish two critical goals: 1-predict emergent phenotypes that stems from the dynamic interplay between G, E and M, and thus enable the breeder to consider the behavior of new genetic and trait combinations in environments that plants have not been exposed or tested before, and 2-identify G x M technologies that could increase food and nutritional security while regenerating natural and production resources. We highlight the need to invest in artificial intelligence and information technologies for breeders to harness multiple sources of information to create G x M technologies to address the diverse cultural and geographically granular societal needs.

## INTRODUCTION

There is consensus that climate change will have a net negative and inequitable impact on agriculture (Lobell et al. 2011; Challinor et al. 2014; Weber et al., 2018; IPCC 2021). The increase in temperature, vapor pressure deficit (VPD) and shifting water balances worldwide will likely change geographical patterns of farming (Ripple et al. 2016; Ficklin & Novick 2017). Disease pressure will increase in high latitudes challenging further crop production and the design of stable agricultural systems (Chaloner et al., 2021). There is agreement that crop improvement will be key to cope with climate change effects on food security (Lou et al., 2009; Cairns et al. 2012; Chapman et al. 2012; Atlin et al. 2017; Hernandez-Ochoa et al., 2019; IPCC 2021; Snowdon et al., 2021; Kholová et al. 2021) and approaches have been proposed (Ceccarelli et al. 2019, Ramirez-Villegas et al. 2015, 2018, 2020, Ceccarelli & Grando 2020). However, if we articulate the problem within a circular economy framework, we can foresee agriculture as part of the solution to climate change rather than the cause of the problem (Bummer et al. 2011; Messina et al., 2022c). Surprisingly, very few efforts were dedicated to answer the question how to breed crops for climate change while regenerating natural resources and reducing greenhouse gas emissions (Brummer et al. 2011, Messina et al. 2022c; Cooper & Messina, 2023).

Historical records indicate that past rate of change in climate was slow relative to the ability of current breeding systems to drive genetic gain as shown for maize in the United States since the onset of a rapid increase in minimum temperatures (Cooper et al., 2014; Messina et al., 2022c). Similar outcomes were achieved for soybeans in the Americas (de Felipe et al. 2016), and wheat despite genotype x environment interactions imposing limits on the rate of genetic gain (Xiong et al. 2021). It is also important to note that until recent decades, rates of genetic gain for maize evaluated under water deficit were lower than under irrigated conditions in the corn-belt of the United States (Cooper et al., 2014; Messina et al., 2022c). The importance of this observation is that only absolute rates of genetic gain are relevant to assess the capacity of the world breeding system to satisfy the demand for food, feed, and renewal fuels (Fisher et al. 2014; Ray et al. 2013). Maize breeding also shows that implementing dedicated breeding programs to improve drought tolerance can deliver germplasm with this characteristic while maintaining yields under water sufficiency in both temperate (Cooper et al., 2014; Gaffney et al., 2015; Messina et al., 2022a) and tropical target environments (Nurmberg et al., 2021; Prasanna et al., 2021). The limited empirical evidence, relative to all crops and all cropping areas in the world, suggests that dedicated breeding efforts proved effective to create adapted germplasm to the target population of environments (TPE) when the mixture and frequency of environment types change at the relatively low pace observed for the past five or more decades.

However, genotype (G) by environment (E) by management (M) interactions (G x E x M) are ubiquitous in agriculture, these are expressed as changes in rankings among genotypes when exposed to different environments and agronomic management practices. Therefore, selecting genotypes for the inadequately defined TPE and/or management inevitable leads to lower than the attainable rates of genetic gain (Cooper & Hammer, 1996; Kholová et al. 2021). There is current evidence that genetic gain in wheat has been hampered by climate change (Morgounov et al. 2013; Xiong et al. 2021). This is particularly critical because whenever the future TPE differs from the current TPE due to climate change, there is a risk that implementing breeding programs for future climates may decrease the current rate of genetic gain for current environments, and thus compromise current food security. There is evidence that wheat breeding may be falling into this trap as rankings of cultivars are changing with climate change (Morgounov et al. 2013; Xiong et al. 2021). Because the traits and trait network interactions underpinning adaptation (at least to drought) change with levels of evapotranspiration (Messina et al., 2011; Borrell et al. 2014; Gleason et al., 2022; Cooper & Messina, 2023), the environmental distance between the current TPE and the future TPE could become significant drivers of G x E x M interactions. In wheat, increased spring maximum temperatures led to both increased and decreased yields (Morgounov et al. 2013). In maize, the conductance response to vapor pressure deficit is a trait underpinning drought adaptation (Shekoofa et al, 2015; Messina et al. 2015). The intensification of drought would magnify the selection pressure applied to the germplasm for elevated levels of limited conductance. The normal operation of the breeding program is expected to be sufficient to increase the frequency of preferred alleles for the required trait levels and thus the breeding program would smoothly match the expression levels of the trait to the changing TPE. In contrast, such as in well-watered dryland systems, the intensification of drought will be conducive to G x E x M interactions and dedicated breeding and agronomy efforts will be required to breed new genotypes for the changing TPE (Messina et al., 2011; Messina et al., 2015; Cooper et al., 2022). The opposite could be said about improvements for radiation use efficiency as evidenced in maize breeding (Messina et al., 2022b). It is anticipated the requirements for the creation of adapted germplasm and cultivars will be inevitably different for different geographies and cropping systems (Kholová et al. 2021). A key question is then *how to harmonize breeding efforts for agricultural systems that regenerate the environment while providing nutrition security to society and improved adaptation to climate change?*

In this chapter we expand on the framework proposed by Cooper & Messina (2023). We use the Breeder’s equation framework and theoretical principles of G x E interactions to demonstrate the critical need to properly time the pace of crop improvement to pace of climate change, and the opportunity to use advanced prediction methodologies to accomplish this goal. Then we articulate the need to rethink breeding objectives to enable agriculture to become part of the solution to climate change (Bummer et al. 2011; Kholová et al. 2021; Messina et al., 2022c). We use an example to demonstrate how to create prediction systems that harness genomic, agronomic, and environmental predictors to implement new breeding objectives within breeding programs and manage and align the breeding program to the changing TPE. We finally discuss the need to enable breeders with dynamic gene-to-phenotype platforms to implement the needed changes in plant breeding and agronomy so society can meet current nutritional and ecosystem regeneration demands, without compromising current and/or future societal needs.

## A FRAMEWORK FOR CROP IMPROVEMENT FOR CLIMATE CHANGE

### Predictive breeding

The goal of a plant breeding program is to create germplasm that solves problems in agriculture and thus create value for producers, consumers, and society (Kholová et al. 2021). Hence, plant breeders define objectives that can seek to improve productivity, quality, and reduce negative environmental footprints, among others. Important breeding objectives in crop improvement programs for row-crop agriculture are yield potential, disease tolerance and yield stability. In fruits and vegetables, breeding objectives also include post-harvest traits, flavor, and appearance among others (Tieman et al. 2017; Collantonio et al. 2022). To achieve these objectives breeders create and evaluate a sample of the germplasm in a sample of the TPE over various stages of evaluation and selection (Fig. 1A). The number of individuals evaluated in fields decreases as the germplasm cohorts advance through the stages of product development and testing, while the number of environments and agronomic management practices sampled increases at the same time (Fig. 1A). By sampling the TPE and the target population of genotypes (Fig. 1B), breeders create prediction models based on statistical and/or dynamic models. The use of managed stress environments helps to expose the germplasm to a set of environments that are of key interest to the breeder (Fig. 1B). These models help them use a relatively small sample of germplasm and environments to explore the complex dimensions of a much larger genotype x environment state space (Fig. 1C; Ramstein et al. 2019). Various genotype-to-phenotype prediction methodologies were developed, some of which are described in this review (Lorenz et al. 2011; Jarquín et al. 2014; Messina et al. 2018; Ramirez Villegas et al., 2020). Plant breeders can use the framework of the “Breeder’s equation” (Lynch & Walsh, 1998) to evaluate genetic progress and optimize breeding systems,

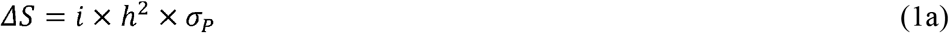

where Δ*S* is response to selection for one cycle of directional selection, *i* is the standardized selection differential applied to a trait, *h^2^* is heritability or fraction of the total phenotypic variation that could be attributed to genetic variation, and *σ_P_* is the is the expected standard deviation for the observed on-farm values of the selection units for the same trait. Prediction methodologies such as whole genome prediction (WGP) seek to use genomic information to maximize the correlation *r_A(M,T)_* between the observed values or breeding values of traits measured on the selection units in the multi-environment trial *M* and the true trait values or breeding values of the selection units in the TPE (Cooper & Messina, 2023),

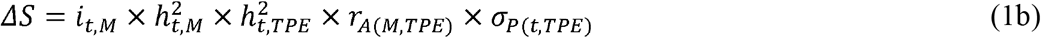

where *i_t,M_* is the standardised selection differential applied to the trait *t* based on the data and/or predictions from models constructed using the sample of environments obtained in trials M, the heritabilities are for the trait estimated for M (*h^2^_t,M_*) and the TPE (*h^2^_t,TPE_*). Prediction models can be based on regression or association within different statistical frameworks (Meuwissen et al. 2001; Yu et al. 2006; Heffner et al. 2009; Zhang et al. 2010; Lorenz et al. 2011),

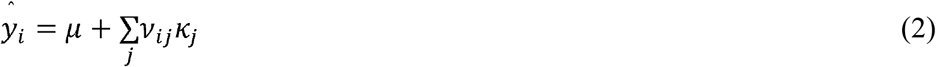

where the trait phenotype *y* for individual *i* is predicted based on the population mean *μ* and the sum over the genotype markers *j* (*ν_ij_*) times the marker effects *κ_j_*. Other models extended this model to include environmental covariates (Boer et al. 2007; Heslot et al. 2014; Jarquin et al. 2014; Millet et al. 2019), consider non-linear associations (Collantonio et al., 2022) or are integrated with dynamical models (Technow et al. 2015; Messina et al. 2018; Diepenbrock et al. 2021). The goal of these prediction methods is to enable breeders to expand their breeding program by adding a virtual component or to maintain the size of the breeding program while using less resources. The efficacy of the application of prediction within large commercial breeding programs for maize has been demonstrated (Cooper et al., 2014; Cooper et al., 2016). Experimental design seeks to reduce experimental error and thus increase *h^2^*, and increase *σ_P_* by exposing the germplasm to environments and management conditions conducive to express variation in adaptive traits in both selection and evaluation environments (Cooper & Hammer, 1996).

**Fig 1.**
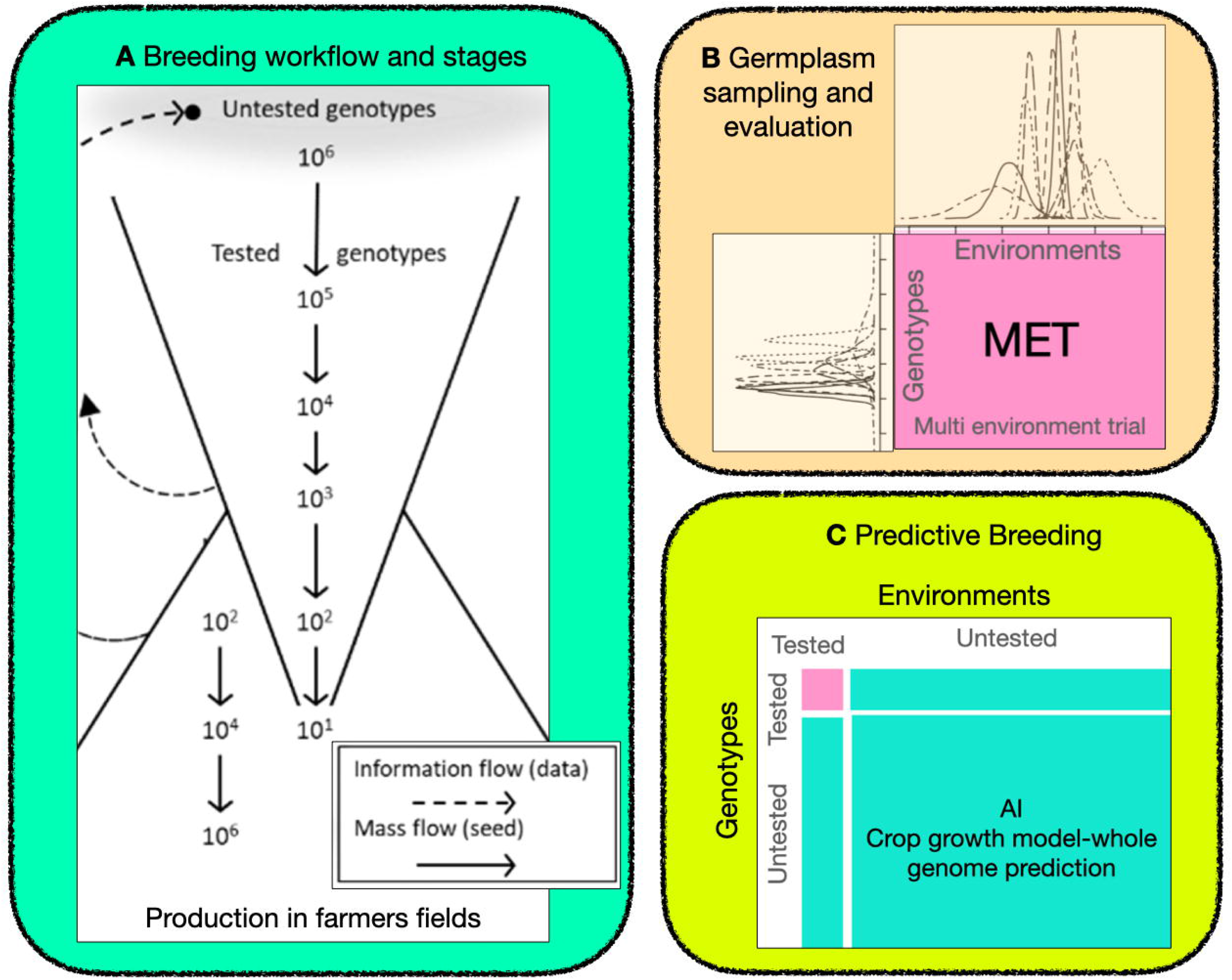
Representation of a breeding program from an operational perspective (A), quantitative genetics view of the sampling of environments within the target population of environments, and sampling genotypes from the target population of genotypes (B), and prediction opportunities that harness the data generated in (B) to augment the size of the breeding program (C) by predicting the possible performance of untested genotypes (A) in untested current and/or future environments (B).

### Genotype x environment interactions hampers genetic gain

Decomposing the heritability into variance components (genetic (G)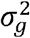; genotype x environment (G x E) 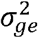; error 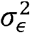; *n* denotes number of environments *e* and replicates *r*) illuminates the potential for G x E interactions to hamper genetic gain (Comstock and Moll 1963, Cooper & Hammer, 1996),

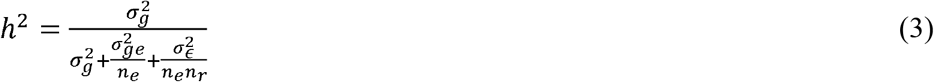

However, the various forms of G x E can have differential impacts on genetic gain (Cooper & Hammer, 1996) depending on the importance of G x E due to heterogeneity of variances (*V*(*σ*_*e*(*env*)_) or lack of correlation. The latter could be decomposed further into *r_g_*, the pooled genetic correlation among environments, and 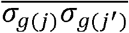, the arithmetic average over all pairwise geometric means among all the genotypic variance for environment *j*’s. Figure 2 shows how lack of correlation can cause changes in the ranking of genotypes while heterogeneity of variances does not. Therefore, to maximize genetic gain it is important to know which environments generate cross over interactions, and what are the frequencies of these environment types within the TPE. In the context of breeding for climate change, it is of critical importance to determine whether future climates will contribute to cross over interactions or not. The encapsulation of crop, soil and environmental science within crop models enables the assessment of traits to understand traits undermining adaptation to current and future climates (Hammer et al., 2014, 2020; Ramirez Villegas et al., 2020; Cooper & Messina, 2023)

**Fig 2.**
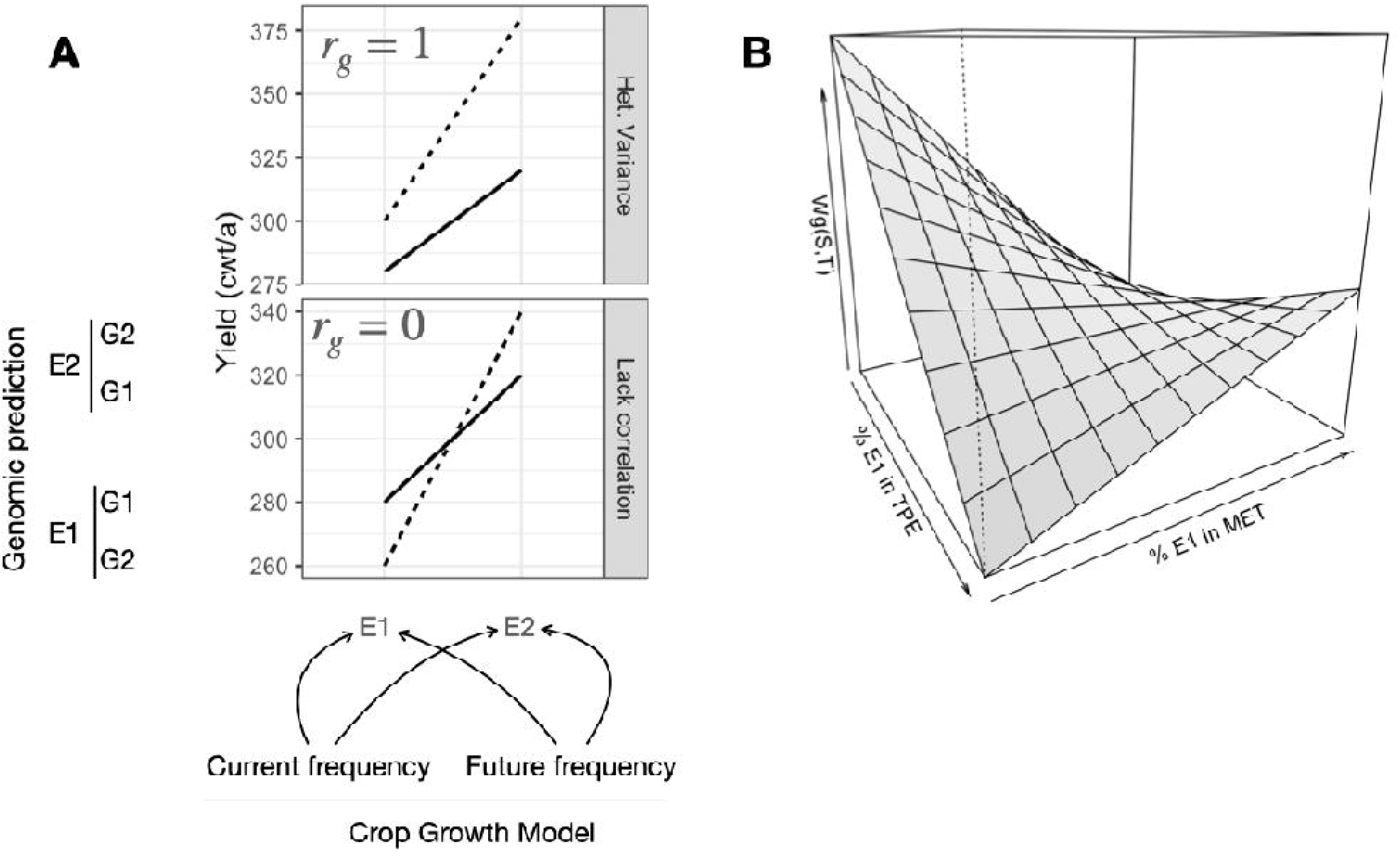
Types of genotype x environment interactions as determined by heterogeneity of variance or lack of correlation (A), and their implication given adequate (inadequate) sampling of environments (%E1) in the target population of environments (TPE) and the multienvironment trail (MET) on the covariance of performance of the selected genotypes (W(S,T), <u>S</u>ample, <u>T</u>arget). Opportunities for the use of crop growth modeling to estimate current and future frequencies of environment types (e.g., E1), and the use of genomic prediction for genotypes G1 and G2, is shown.

### Environment frequencies and weighted selection

Because of the ubiquity of G x E interactions, and the possibility of biases in the sampling of environments by the implemented testing system within the breeding program, selection strategies that accommodate G x E interactions were developed (Cooper et al. 1995), and weighted selection strategies have been proposed (Podlich et al. 1999). In this case, weights are estimated from the relative frequency of sampled environments over the expected frequencies of the environment types that comprise the mixture of environments of the TPE. Frequency of environments could be defined based on climatology or more sophisticated methods (Chapman et al. 2000; Löffler et al. 2005; Kholová et al. 2013; Ramirez Villegas et al., 2020; Cooper & Messina, 2021; Carcedo et al. 2022), such as crop growth models (CGM, Jones et al. 2003; Holsworth et al., 2014). CGMs are functions that approximate the phenotypic function,

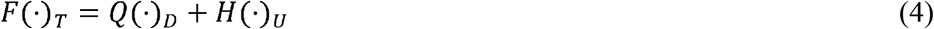

where the function *F* represents perfect knowledge of the observable phenotype as determined by nature, the function *Q* represents the phenotypes that are predictable based on our current scientific knowledge and phenotyping systems, and *H* is the function that represents what we don’t know, it is not knowable/predictable (Day 1976), or we do not want to include in the model. If we define function *Q* as a CGM, the CGM becomes a cognitive construct that we use to represent processes, states, and the topology of relations among biological processes (Cooper et al., 2009). CGMs can span several levels of organization, however, there is consensus that empiricisms are needed whenever we seek to link or use information for more than two levels of organization away from the target level of the system to be modelled (e.g., cell to crop). CGMs have a set of parameters that are known *(k)*, unknown and estimable *(u)* and a set of inputs (*I*), *CGM*(*x_k_*,*x_u_*, *I*). Using soil, weather and management databases as inputs, Harrison et al. (2014) estimated frequencies of environment types for current and future climates. This knowledge could enable breeders to implement breeding strategies that create a smooth transition from current to future genotypes adapted to climate change, that accompanies the changes in the mixture of environments in the TPE. Using genomic predictions in the form of eq (2), and weighted selection will be most useful whenever lack of correlation G x E is an important determinant of future yields.

Figure 2b shows a theoretical representation of the plausible consequences of inadequate decisions regarding selection for climate change in the presence of both types of G x E interaction (Fig. 2A). Taking E1 as current climates, the percent frequency of E1 in the TPE indicates the transition from current to future climates. When the frequency of environment type E1 matches the frequency in the TPE, say evaluation in the current climates for selection in the current climates, the covariance *W*(S,T) between selection (S) and target (T) is highest. The difference at the extremes is due to the heterogeneity in the variance in E1 and E2 (Fig. 2A). Because of the lack of correlation between environment types, selecting for E1 when the target is E2, that is selecting in current environments for performance in future environments, can lead to negative covariances. Most of the arguments to start breeding for climate change are based on this concept (Cairns et al., 2012; Atlin et al., 2017). Lack of action can lead to global food insecurities and geographical famines with consequent starvation of future human populations; there is a merit in this argument because of the ubiquitous G x E in major cropping systems subject to drought stress (Cooper, 1999; Messina et al., 2015; Xiong et al., 2021). However, the opposite is also possible, and premature selection for future climates (E2) can hamper genetic progress for current climates contributing to food insecurity for current populations. This analysis leads to the following propositions:

- it is critical to time correctly breeding strategies with expected changes in climate,
- computational methods in with genomic prediction could be used in combination with weighted selection and crop models to create cultivars and germplasm for a dynamic TPE
- predictive breeding connected with climate predictions can time the required adjustments within breeding programs correctly to produce cultivars ready for use by famers in alignment with changes in the TPE (Challinor et al. 2016).

Because breeding is a dynamic process, careful decisions need to be made and the evolution of the germplasm monitored. Simulation studies have demonstrated that breeding programs are sensitive to the founding germplasm used to start the breeding program (initial conditions) and can manifest temporal patterns that could diverge with increasing cycles of selection (Messina et al., 2011), in similar ways as the weather and climate systems do (IPCC, 2021). The prediction of the plausible trajectories of breeding programs seeking to create the germplasm required for adapting agricultural systems to climate change adds another layer of complexity and uncertainty. This requires careful consideration of breeding objectives for climate change.

### Rethinking breeding objectives

In face of climate change and the need to regenerate natural resources, we advocate for a new framework in which breeding objectives are defined by answering the question: how to use genetic and agronomic levers together to maximize the societal benefit of a unit of resource use (Cooper et al., 2020; Hunt et al. 2019, 2021; Zhao et al. 2022), and how to minimize environmental degradation (Brummer et al. 2011; Rodell et al. 2019; Messina et al., 2022a,c)? Reimagining breeding to enable cropping systems that can improve the efficiency of water use (Blum, 2009; Cooper et al. 2022) is paramount in the context of the temporal changes in patterns of seasonal water availability for food production worldwide (Rosegrant et al. 2009; Richey et al. 2015; Rodell et al, 2018; Caparas et al. 2021). Similarly, given the importance of nitrogen in food production, nitrogen oxides emissions from agricultural fields on global warming, and nitrates on water pollution (Roberston & Vitousek, 2009; Boules et al., 2018; Chang et al. 2021), it is imperative to articulate breeding objectives to create germplasm and cultivars that enable the creation of systems with high efficiency of nitrogen use, and low nitrogen oxides emissions.

Dynamic CGMs are non-linear functions of environment and management inputs, and genetic parameters that simulate with time steps from hourly to daily, the soil carbon, nitrogen and water balance, and the plant carbon and nitrogen balance (Jones et al. 2003; Holsworth et al. 2014). Thus, they would enable breeders to incorporate effective water use, nitrogen losses and emissions, soil carbon and other metrics as components of the breeding objectives. Unlike the past, breeders would have the information needed to create cultivars to minimize externalities, maximize effective water and nitrogen use, soil carbon accumulation, or a combination that contributes to produce food while combating climate change (Messina et al. 2022c; Cooper & Messina, 2023). While improving confidence in the models to simulate the metrics of interest is an important and likely costly undertaking, these investments pale relative to the massive socioeconomic consequences of no action. Traits that contribute to the improvement of both productivity and reduced externalities need not be directly related to the biological efficiency of resource use, as shown for nitrogen (Muller et al., 2019) and water in maize (Cooper et al. 2014; Messina et al. 2022a), but at the cropping system level. It has been proposed there is a need to increase the circularity of nutrient cycling in maize by improving maize tolerance to low temperatures and remobilization to roots (Buckler, pers. comm., Fig. 3). Figure 3 illustrates a concept where tolerance to cold stress in maize could increase yields, synchronize soil nitrogen supply with crop demand, increase light interception, and thus reduce externalities while increasing productivity. Such system level thinking enabled by CGMs could be conducive to a shift in mindsets on how to define breeding objectives that move us away from a plant/crop centric idiotypes with the focus on productivity (Perego et al. 2014; Rötter et al. 2015; Paleri et al. 2017; Hammer et al. 2020; Ramirez Villegas et al., 2020) towards a system centric thinking with a focus on the balance between providing nutrient security while minimizing environmental externalities and their negative contributions that hasten climate change; these are two important dimensions contributing to current and future human health and well-being.

**Fig. 3.**
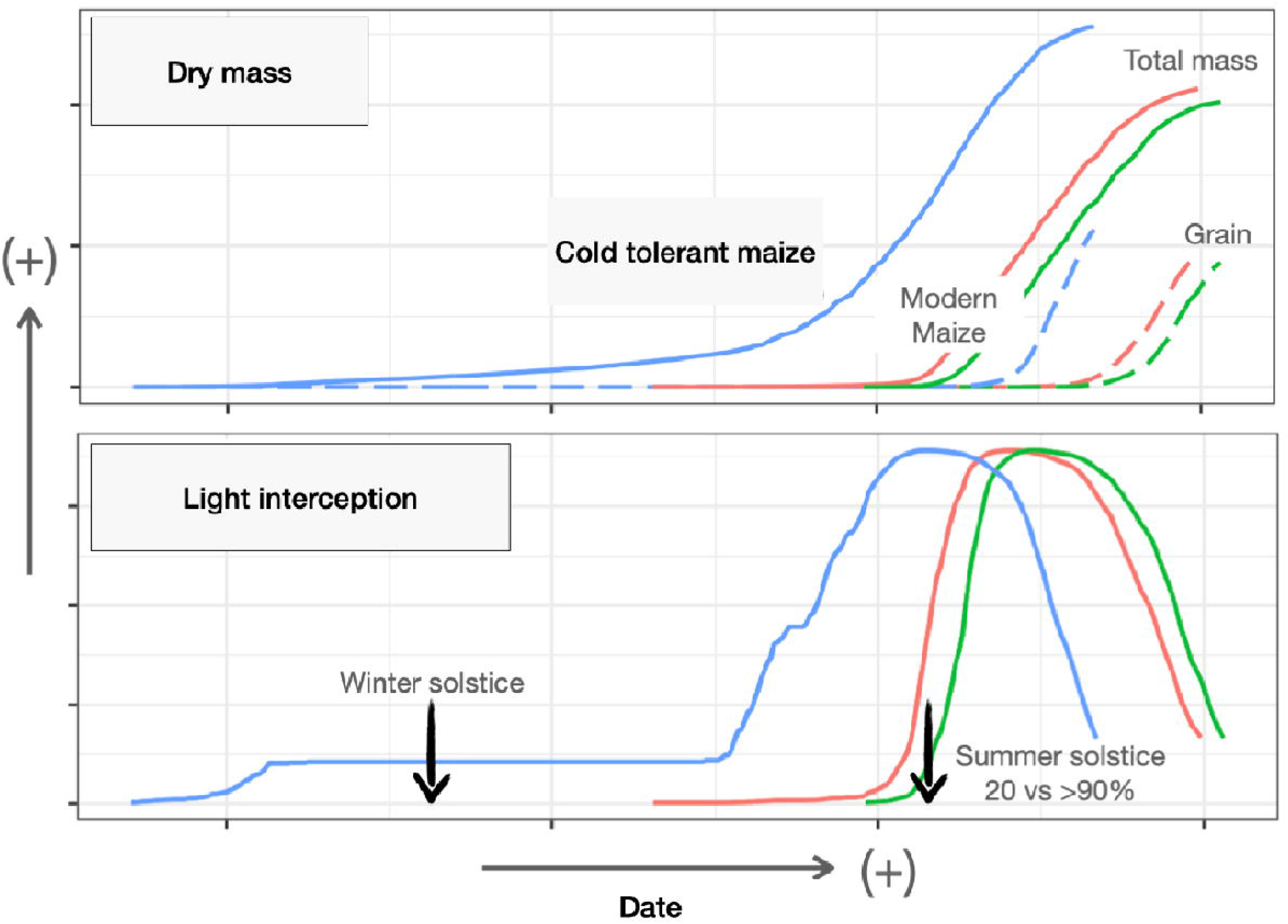
Dry mass growth (total and grain) and light interception for modern maize planted at normal and early planting, and for a conceptual novel maize genotype tolerance to cold. Cold tolerant maize could enable higher yields with lower nitrogen (N) inputs due to improved fertilizer recovery efficiency, timing between N supply and demand, and circularization of N within the system.

### Models that predict emergent phenotypes

For a CGM to deliver on the promise to add information to re-think breeding objectives and predict with higher skill beyond the current sample of environments, they must be able to predict what we define as emergent phenotypes. These are observable phenotypes that are different from what we can predict by understanding the parts in isolation or as independent components (Anderson, 1976; Roeder et al., 2021; Powell et al., 2022). It has been shown that simple equations can have complex behavior, for example the logistic equations that model populations of plants can manifest complex emergent behaviors such as bifurcations and chaos (May, 1976; Roeder et al., 2021) that conform well with the observation of bimodal distributions of maize barren plants (Edmeades & Daynard, 1979). Routines recently implemented in the crop model APSIM can simulate emergent patterns such as the relationship between anthesis-sinking interval and yield in maize or the relationship between plant growth and kernel numbers (Messina et al. 2019). While the routines in the current APSIM model do not yet have the capacity to simulate the observed bifurcation behavior of barren and non-barren plants in a crop, the capacity to predict emergent phenotypes suggests that dynamical models are tools that are evolving and are capable of predicting phenotypes different from what is expected from the analyses of the parts in isolation. For example, understanding a system to reduce photorespiration in leaves in isolation leads to an overestimate of the plausible impacts on productivity at the crop scale and for the mixture of environment types that comprise the TPE (Hammer et al., 2019). This prediction using APSIM conforms well with the limited success of large numbers of single gene transformation technologies for yield improvement (Simmons et al., 2021). The ability of current CGMs to simulate some emergent behaviors, can enable breeders to use this kind of tool to inform selections to improve germplasm to meet the needs of the adjacent environment space of the future TPE while considering the expected change in frequencies of environments in the selection and on-farm production situations (Snowdon et al. 2021; Messina et al. 2022a,c). Because the CGM integrates scientific knowledge, for many situations these tools can be better equipped than purely empirical models to predict genotype performance in environments that were never included in the training sets, and more importantly, environments that the crops have never experience before (Battiest & Naylor, 2009).

CGMs are also useful tools to explore the relevance of traits and trait networks (Messina et al. 2020; Ramirez Villegas et al., 2020; Gleason et al. 2022), and genetic networks (Messina et al. 2011; Powell et al. 2022) on yield performance. Figure 4A shows the contribution of individual traits or trait networks (TxT) to the fraction of phenotypic variance. At the environment extremes of high and low evapotranspiration (ET), trait interactions are less relevant with either growth traits or reproductive resiliency traits explaining most of the variation in this maize example. At intermediate levels of ET, the interaction of traits as trait networks underpins most of the phenotypic variation. In the presence of these trait interactions and interactions with the environment, we have observed that a CGM can perform better than empirical models (Technow et al. 2015, Messina et al. 2018, Diepenbrock et al. 2022, Messina et al. 2022c). Cooper et al. (2009) using simulation approaches argued that the benefit of a molecular breeding strategy over phenotypic selection increases with increasing complexity of the genotype and environment system. Diepenbrock et at (2022) showed that improvements in predictive skill by incorporating biological knowledge to the prediction algorithm increase with decreasing ET (Fig. 4B) in agreement with theoretical predictions (Cooper et al., 2005; Cooper et al., 2009; Messina et al., 2011). The emergence of G x E interactions also depends on the distance between environments (E1 to E6, Fig. 4). In figure 4A we illustrate, based on the tradeoffs determined by biophysics embedded in the CGM, various types of G x E interactions. If climate change determines a rapid shift in ET form E2 to E1 (CC2→1), we should not expect major changes in G x E and thus the normal operation of a breeding program will suffice to adapt the germplasm to the changing TPE; physiological basis of adaptation moves from complex (high fraction of the variance explained by trait x trait interactions) to simple (few interactions underpin adaptation). At contrast, if climate change determines a shift from E5 to E3 (CC5→3), a drastic increase in G x E interactions can emerge underpinned by new trait x trait interactions, and deliberate breeding augmented by CGM, and genomic prediction can be expected to enhance the potential to harmonize improved germplasm adaptation with the rate of environmental change. Yields in E5 are largely determined by growth (e.g., RUE, Messina et al. 2022b) that requires large quantities of water. Genotypes with high RUE that will rank high in E5, will rapidly consume soil water and yield poorly in E3 under severe water deficits. Similar examples could be drawn for soybean (Sinclair, 2011), chickpea (Sivasakthi et al. 2017) and other crops. Traits such as root angle can also affect the dynamics of water use, and thus the expression of emergent phenotypes and trait interactions; genomic regions associated with root angle were implicated in the maintenance of green canopies and thus biomass assimilation through photosynthesis post-flowering (Manschadi et al. 2008; Mace et al. 2012). In E1 environments, reproductive resilience and not water capture or pattern of water use is the main determinant of yield under stress (Messina et al. 2021).

**Fig 4.**
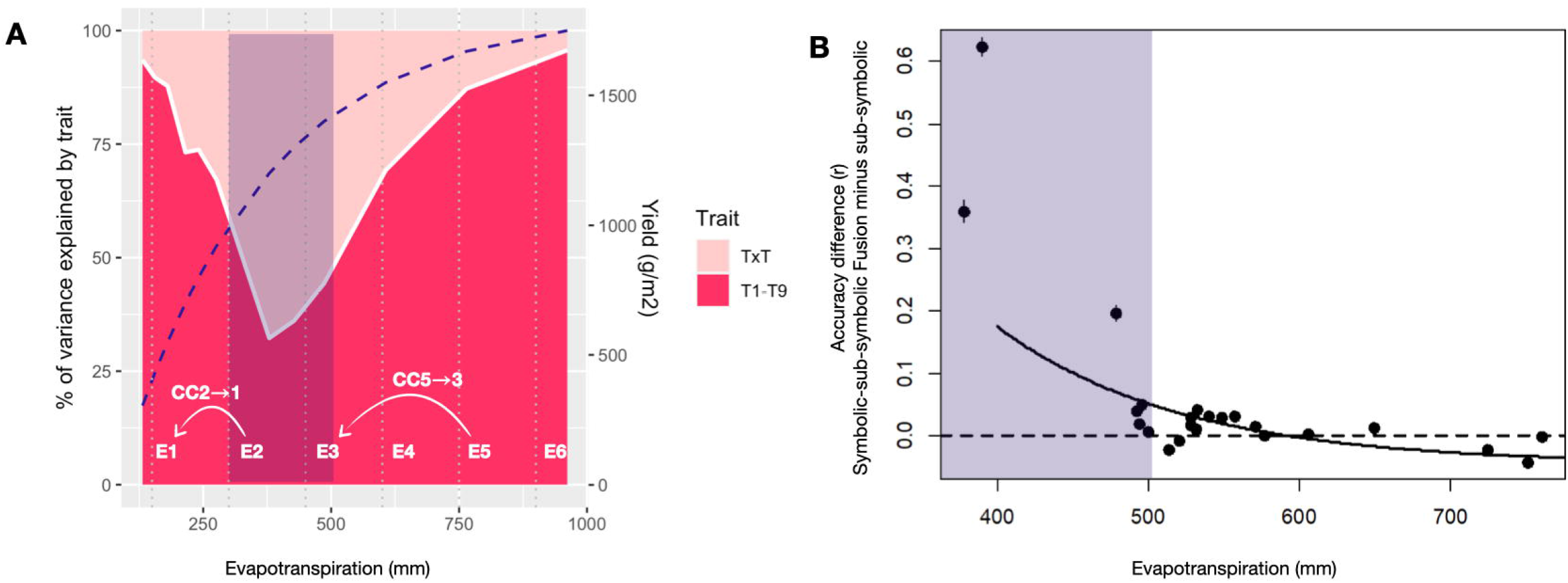
Percent of variance in maize yield explained by the sum of traits or trait interactions (A) and accuracy difference (correlation coefficient, r) between a symbolic-sub symbolic fusion approach (crop model + genomic selection) minus sub-symbolic approach (genomic selection) along an evapotranspiration gradient (B). Purple areas in A and B highlight the increase in prediction accuracy difference between methods correlates with the importance of trait x trait interactions on the determination of yields. Environment E1-6 shown to illustrate plausible examples of changing in climate (CC5➔3; CC2➔1).

### Example: Integrating environmental and genomic predictions using crop models

Equation 2 corresponds to a simple linear model for a genomic predictor, this could be yield across environments or more than one environment type. While we often think about 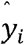 as yield for genotype *i*, breeders often predict other traits such as flowering time, time to maturity or grain moisture at maturity. Objective and/or subjective indices are used to integrate these predictions for multiple adapted traits. Millet et al. (2019) used various approaches to predict yield and yield components and were able to build a static crop model and use environmental covariates to predict some important GxE interactions for yield. These static crop models are often linear, and unable to generate unexpected or emergent behavior that may surface when predicting new genotypes in new environments, that is into a G x E space that was not included in the training set or is beyond the environments that the germplasm was generally grown (Battiest & Naylor, 2009; Hammer et al., 2019). Recall, that the relation between growth and kernel number in maize is non-linear (Andrade et al., 1999), so linear approximations are limited from their conception. However, the same linear static model applied to the prediction of the rates of change in physiological processes in response to environmental variation can lead to emergent patterns of G x E upon numerical integration over the growing season (Messina et al. 2019; Hammer et al., 2019) and over cycles of selection (Chapman et al. 2003; Messina et al. 2011). This approach was demonstrated in various crops including soybean (Messina et al. 2006), dry bean (Hoogenboom et al. 2004), Maize (Messina et al. 2011), sorghum (Chapman et al. 2003), and Barley (Yin et al. 2003) among others. The experimental demands often limit the applicability of these models when many loci underpin the control the traits of interest.

Bayesian approaches were proposed to overcome this limitation, estimate genetic and physiological models simultaneously, and deal explicitly with the levels of uncertainty,

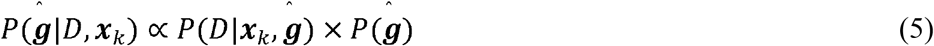

where 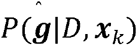 is the posterior distribution, conditional to the vector of traits that were kept constant (**x**_*k*_), 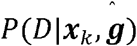 is the likelihood function with data (*D*) generated by the crop models with **x**_*k*_ known or estimable parameters (marker effects, 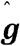), and 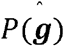 is the prior distribution of the traits and the marker effects for the traits. Figure 5 shows a diagram for the estimation of a single trait, RUE, which it is adapted from Technow et al. (2015) and implemented as an Approximate Bayesian Computation approach. The procedure consists in sampling the posterior distribution of RUE to assign values to the various genotypes in the training set. A genomic predictor for RUE based on markers (eq. 2) is estimated; Genomic Best Linear Predictors (GBLUPs) are generated for each genotype. Phenotypes for which the breeder collected data (yield, flowering time, kernel numbers, etc) are predicted using the CGM with environmental inputs for the trial and model parameter/s estimated using genomic predictors such as GBLUP for the physiological trait, here RUE. The distance between the observed and the predicted phenotypes is calculated. Using a metric such as root mean square error, others are also valid for individual traits or trait combinations, a decision is made based on a rejection algorithm to keep the sample and build the posterior distribution.

**Fig. 5.**
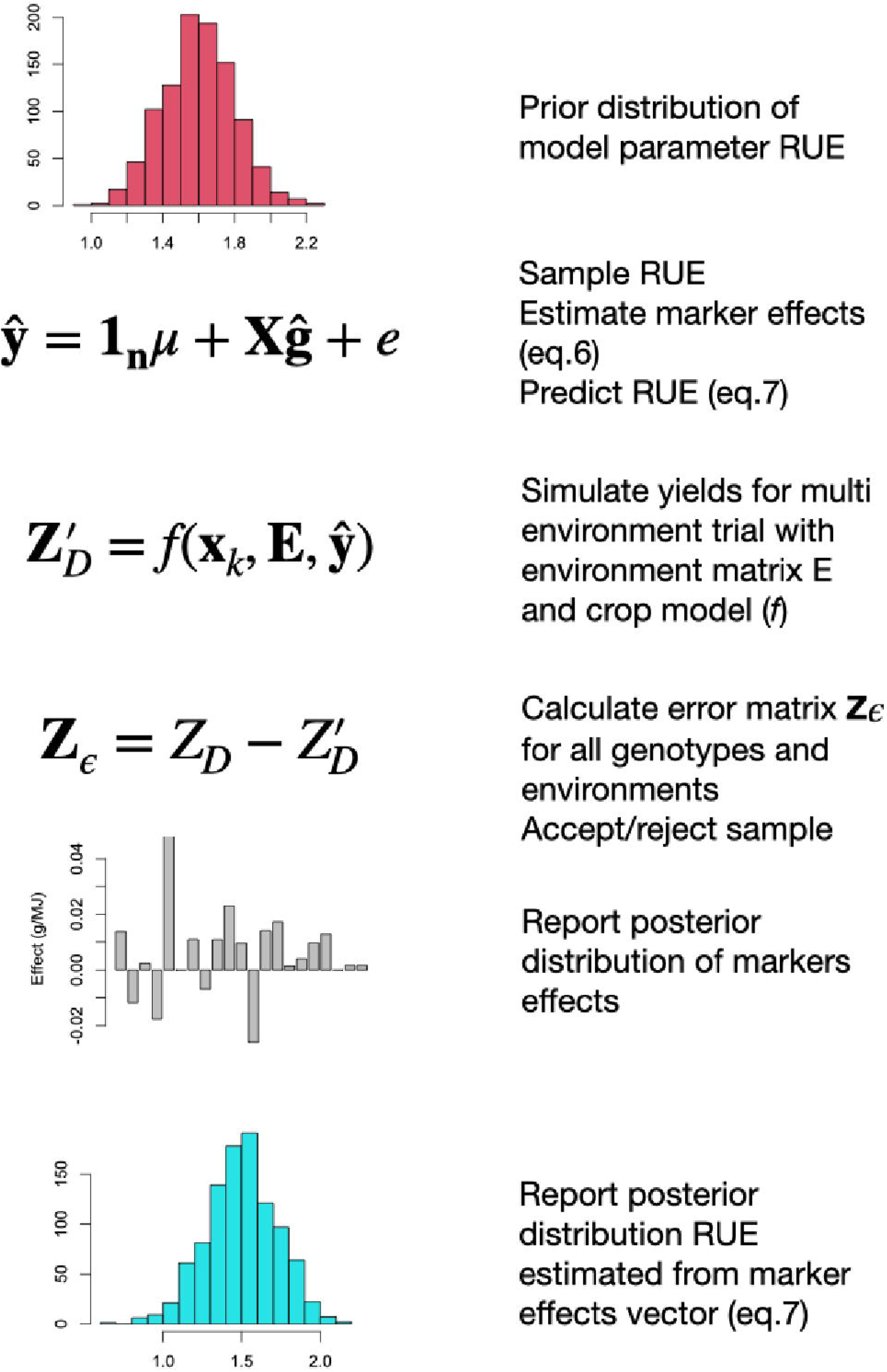
Pseudo algorithm to describe the workflow of a crop growth model – genomic selection approach for a single crop and a single trait (radiation use efficiency, RUE) that uses genomic best linear predictors at the trait level **y** from makers (**X**), marker effects **g**, environment inputs **E**, and set of fixed crop model parameters **x**_k_.

In the example presented in figure 5, the mean 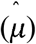 and vector of marker effects for RUE were modeled as,

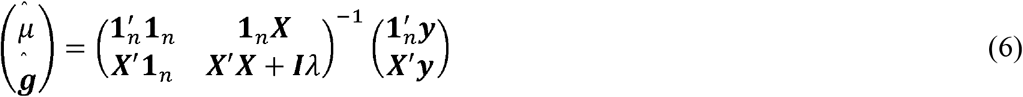

where *X_nxm_* is a matrix of *n* genotypes by *m* molecular markers, **y** is a vector of RUE values, and *λ* is a parameter to describe the signal to noise ratio in the data. The RUE for each genotype is estimated from the molecular markers as,

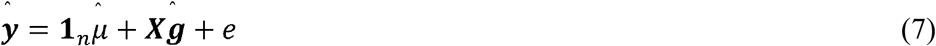

The posterior distribution was built from marker vectors for which the root mean square error for the yield prediction was less or equal to 20% of the mean. Pseudo-code for this algorithm is presented in Technow et al. (2015).

Figure 6 shows the simulation experiment whereby a sample of genotypes for various levels of RUE was generated. The map between RUE and simulated yields for two environments contrasting for water availability is shown (Fig. 6A). The bifurcation is generated by how the model simulated the pattern of water use, and thus how much water was allocated to vegetative growth vs. reproductive growth. In the water sufficient environment E1 (Fig. 6A), yield increased with increasing RUE. The opposite occurs in E2 where high RUE leads to high water use pre-flowering and induces drought during grain-fill (Fig. 6A). This result agrees with experimental observations (Cooper et al. 2014), large scale simulation for the limited transpiration trait in maize in the U.S. (Messina et al. 2015) and yield data (Adee et al. 2016). When RUE is predicted using markers the predicted range of RUE is lower than the observation, as expected from the use of the GBLUP method that shrinks predictions towards the mean in proportion to the signal to noise ratio. However, the CGM linked to a whole genome prediction method such as GBLUP (CGM-WGP) can regenerate the emergent relation between RUE and yield (Fig. 6B). Messina et al. (2018, 2022c) and Diepenbrock et al. (2022) used a Metropolis-Hasting within Gibbs algorithm and showed that a) the advantage in prediction accuracy of CGM-WGP over WGP (Bayes A) for yield increases with increasing complexity of the target environment quantified by decreasing ET, b) the use of multiple traits can further increase predictive skill of CGM-WGP, and c) the CGM-WGP can borrow information by using the trait relations embedded in the CGM to make predictions on phenotypes that were not measured or used in the training of the model. The ability of CGM-WGP to predict time to silking and kernel numbers in the absence of silking and kernel number data (Messina et al. 2022c), when the model was trained on yield alone, is a promising result to encourage further development of approaches to predict other state variables that are more difficult to measure such as effective water use, soil nitrogen losses, soil water recharge, and soil carbon accumulation.

**Fig. 6.**
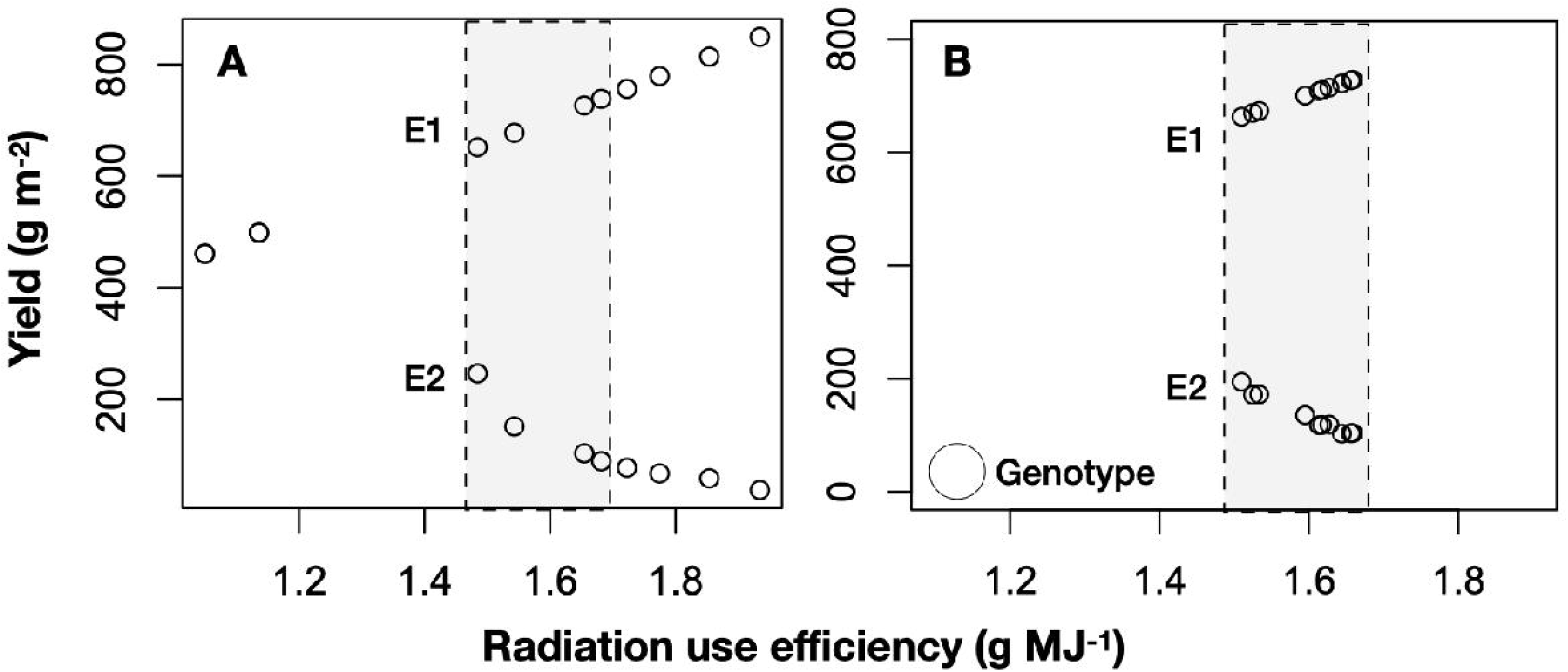
Simulated (A) and predicted using crop model-genomic selection (B) yield response to radiation use efficiency for ten genotypes (circles) in two environments (E1, E2)

### Enabling breeders with dynamic gene-to-phenotype platforms

The research conducted over two decades in maize in the U.S. (Messina et al. 2020; Cooper & Messina, 2023) is informative to consider what the scientific community and breeders working in other crops and geographies can expect. Our research studying how predictive skill differentials between pure statistical approaches compared with those augmented by dynamical models (Diepenbrock et al. 2022), and how trait networks and trait interactions underpins the variance explained by these components (Messina et al. 2020, Gleason et al. 2022; Cooper & Messina 2023) indicates that CGM-WGP approaches can be expected to have an increasingly important role in breeding for the impacts of climate change. The exacerbation of climate extremes within the TPE, because of a changing climate (IPCC, 2021), will be conducive to increasing the distance between environment types encountered within the TPE. In figure 6 we show how different traits underpin adaptation to these extreme environment-types and how these traits can lead to the emergence of lack of correlation G x E interactions (Fig. 2), and thus an increasing challenge to maintain or increase rates of genetic gain within the TPE. Moreover, the results presented in Figures 2 and 4 suggest that the strategies for improving crop adaptation will be regional in nature and the climate change induced distances among environment types and their frequency of occurrence are expected to vary across geographies.

The development and deployment of platforms capable of predicting genotype x environment x management interactions (APSIM, Holzworth et al. 2014; DSSAT, Jones et al., 2003) by harnessing environmental and genomic predictors within a physiological framework (e.g., Peng et al. 2020; Diepenbrock et al., 2022) for many different crops and geographies is a research imperative in the years to come. Evaluating and improving CGM-WGP will require engaging breeders, agronomists, and physiologists/modelers, who can expand the CGM-WGP capability beyond maize and to make it accessible to the broader community. Albeit limited, the evidence available to date suggests that yield-trait performance landscapes for agricultural systems are complex (Chapman et al. 2003; Messina et al. 2011; Hammer et al. 2006, 2014, 2020) and there will be a need for resources and information technologies to enable simulation at very large scale to enable the world community of breeders to explore and leverage the knowledge hidden within a massive gene-to-phenotype spaces (Cooper et al., 2009; Ramstein et al. 2019; Cooper et al., 2020; Cooper et al. 2022).

CGM-WGP technology on its own will be limited in the type of solutions it can bring to address the climate change challenge. Gap analyses, the study of actual crop yields in farmer’s fields in the context of resource use and attainable yields for that level of resource use (Lobell et al. 2009; van Ittersum et al 2013; Cooper et al., 2020) provide a framework within which the breeder and the agronomist can search the G x E x M state-space and develop G x M technologies for the patterns of change they expect in the TPE as a consequence and in line with the rate of change due to climate change; G x M technologies that can close the yield gap taking into consideration the expected frequencies of E environment-types in the current and future climate-affected TPE. This plant breeding-agronomy gap analyses framework could be a foundational method to translate CGM-WGP technology into practical G x M technology applications by answering the questions posed by breeders (what are the best genotypes for the TPE), the agronomists (which genotype subset fits best the cropping systems in my geography, which sets of genotypes would enable me to innovate at the cropping system level?), and the farmer question, which genotypes perform in my operation with the resources available to me for production? (Fig. 7). Answering these questions will require harnessing AI, involve transdisciplinary thinking, and embrace circularity in agricultural production.

**Fig 7.**
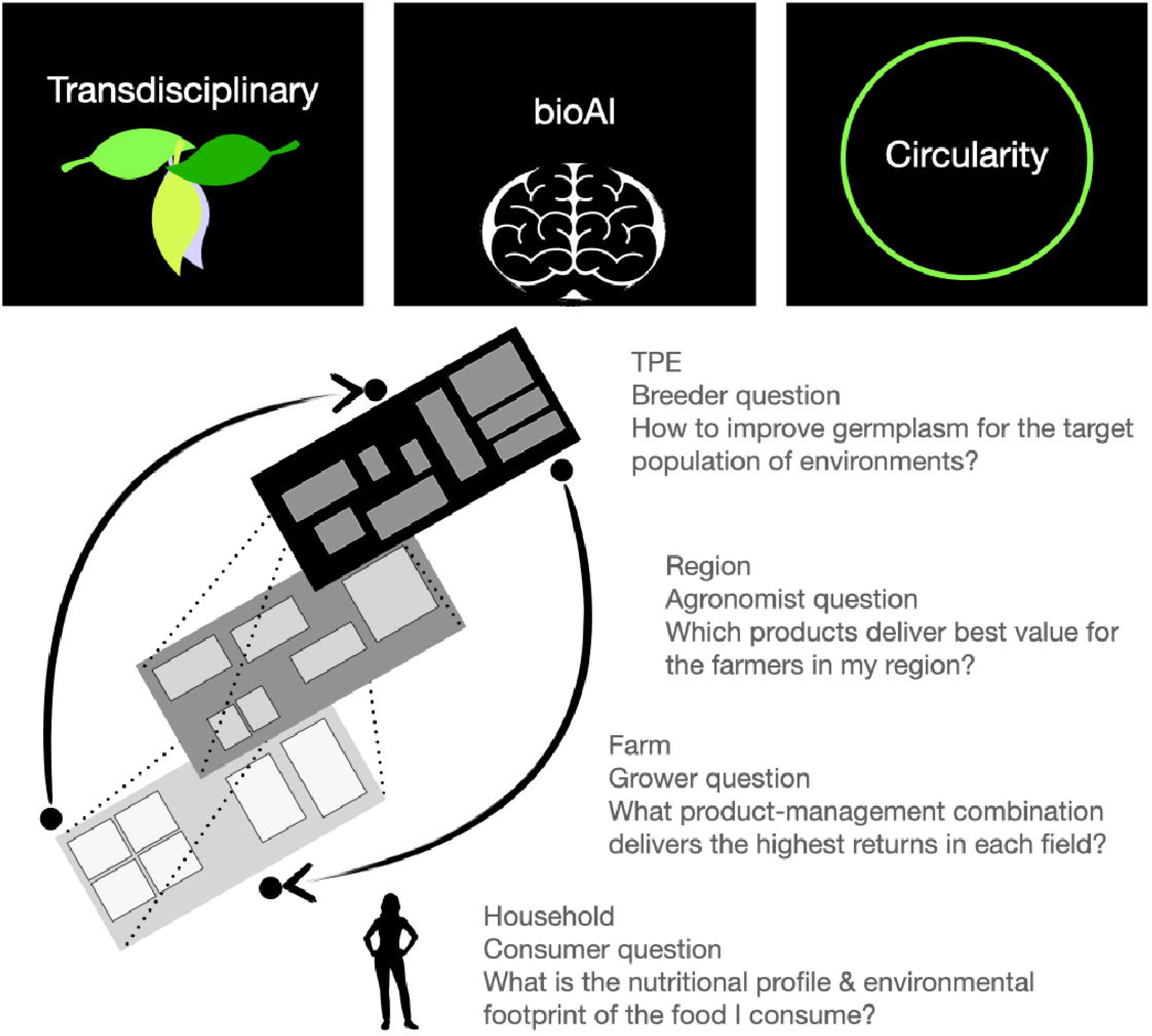
Transdisciplinary teams will be required to keep a dialogue going to answer questions from breeder to household, which answers will enable society to create systems adapted to climate change. Given the complexity of the methodologies that will be required to identify and design genotype, management, and agricultural systems, we foresee the need to develop a biologically grounded symbolic and sub-symbolic integration of artificial intelligence methods (bioAI). Principles of circular economies would enable the design of agricultural systems that could be part of the solution to climate change.

## 6. PERSPECTIVES

Genetics for crop improvement ranks in the top set of technologies that can contribute to human adaptation to climate change. Plant breeding has contributed to yield improvement since the domestication of crop species, and it was most evident in the last century. In this chapter, we have shown the need to develop new G x M technologies to adapt to climate change, but also, the need to transition from current to future G x M technologies in a way that future food security does not come at the expense of current food security. To spur scientific and technological development for climate change adaptation, we have demonstrated an approach to integrate genomic, agronomic, and environmental predictors to accomplish three critical goals

- predict emergent phenotypes that stems from the dynamic interplay between G, E and M, and thus enable the breeder to consider the behavior of genetic and trait combinations in environments that plants have not been exposed to or tested within before, and/or in more variable and extreme environments,
- identify G x M technologies that could increase food and nutritional security while regenerating natural and production resources; outputs from crop model-based approaches could be used to create multi-dimensional selection indices,
- enable breeders and agronomists with tools to harness multiple sources of information to create G x M technologies to address the diverse cultural and geographically granular societal needs.

Breeding can enable incremental adaptation by improving upon existing crops, but also transformational adaptation by enabling new farming systems and new crops. The same toolset, approaches and needs introduced in this chapter for single crops, could be used by a teams focused on crop improvement for new agricultural systems. Soybean-maize production systems like Safrinha in Brazil took forty years of research in breeding and agronomy. Society can no longer afford long time-lags to develop sustainable technologies to adapt cropping systems to climate change. The implementation of dynamic gene-to-phenotype platforms, such as CGM-

WGP within farming systems simulators such as DSSAT and APSIM seems a natural next development to shorten the development time for discovery, refinement, and optimization of novel agricultural systems. To effectively search the extremely large search spaces, now G is expanded to G within crop within a crop sequence, investment will be required in both artificial intelligence and information technologies, to effectively search these spaces, manage very large data sets and run the simulations at unprecedented large scale.

As society continues to make environmentally and health-conscious decisions and increase the demand and consumption of plant-based foods in diets, breeding objectives will continue to evolve to deliver more nutritional value over quantities and reduce environmental footprints. Expansion will be needed to hasten breeding in fruits, vegetables, and pulses, which can benefit from applying breeding technologies previously primarily developed for row crops. The framework introduced in this chapter, based on CGM-WGP or more generally AI could have the largest impacts in breeding programs with small footprints by maximizing information extraction from data and enabling prediction for future environments.

